# Characterization of alkaline stress tolerance mechanisms in *Lotus* forage species modulated by *Pantoea eucalypti*

**DOI:** 10.1101/2020.06.06.138230

**Authors:** Maria Paula Campestre, Nazareno Luis Castagno, Cristian Javier Antonelli, Vanina Giselle Maguire, Francisco Jose Escaray, Oscar Adolfo Ruiz

**Affiliations:** Instituto Tecnológico de Chascomús (INTECh). UNSAM-CONICET. Chascomús, Buenos Aires, Argentina

**Author notes:** **Abbreviations** NPQ, non-photochemical fluorescence quenching; PSII, photosystem II; q_P_, photochemical fluorescence quenching coefficients; ROS, reactive oxygen species; ◻_PSII_, quantum efficiency of PSII in light; Lt, Lotus tenuis; Lc, Lotus corniculatus; Lt × Lc, Interspecific accession Lotus tenuis × Lotus corniculatus.

**Keywords:** *Lotus* spp., Interspecific hybridization, Fe^2+^ homeostasis, Alkaline tolerance

## Abstract

This study was designed to elucidate the physiological responses of three *Lotus* forage accessions to alkaline stress and the influence of the inoculation of a *Pantoea eucalypti* endophyte strain on its mitigation. One-month-old diploid accessions of *Lotus corniculatus* (Lc) and *Lotus tenuis* (Lt), and the interspecific hybrid LtxLc obtained from these parental accessions, were exposed to alkaline stress (pH 8.2) by the addition of NaHCO_3_ 10 mM to the nutrient solution for 2 weeks. The results indicated that Lt and the LtxLc hybrid are alkaline-tolerant compared to Lc, based on the observation that their dry mass is not reduced under stress, and symptoms of chlorosis do not appear on leaf blades, in contrast to observations of the Lc accession subjected to identical growth and stress conditions. In Lc and LtxLc accessions, the Fe^2+^ concentration decreased in the aerial part under stress and increased in the roots. Interveinal chlorosis observed in the youngest leaves of Lc during alkaline treatment was accompanied with a higher reduction of Fe^2+^ levels in shoots and a higher increment of Fe^2+^ in roots, compared to the other accession. Plant inoculation also tended to acidify the medium under alkalinity, contributing to Fe accumulation in the roots. Moreover, the inoculation caused a considerable increase in Fe^2+^ content in shoots in all three *Lotus* forage species under alkaline treatment.

F_v_/F_m_ and PI_ABS_ were only reduced in Lc under alkaline treatment. Inoculation reverted this effect and improved the ABS/RC and DI_o_/RC ratios in all three accessions. In addition, under alkaline conditions, Lc dissipated more energy than control plants. Expression of the metal-transporting gene NRAMP1 increased in the inoculated Lc accession under stress, while remaining unmodified in Lt and LtxLc hybrid.

Altogether, the results obtained make clear the importance of inoculation with *P. eucalypti*, which contributed significantly to the mitigation of alkaline stress. Thus, all the results provide useful information for improving alkaline tolerance traits in *Lotus* forage species and their interspecific hybrids.

## INTRODUCTION

The genus *Lotus* (Fabaceae) includes over 100 forage species and has worldwide distribution [1]. Most species are native to Europe, Asia, Africa and Australia, and some to the Atlantic and Pacific Ocean Islands [2]. The main forage species of genus *Lotus* cultivated in South America are birdsfoot trefoil (*Lotus corniculatus* L.) (Lc) and narrow-leaf birdsfoot trefoil (*L. tenuis* Waldst & Kit) (Lt). These legumes vary in their ability to grow in low fertility soils and under different unfavorable environments [1,2]. While Lt grows well in limiting soils conditions, commercial cultivars of Lc need soils with high or moderate fertility [3].

The Flooding Pampa (Argentina) is a typical cattle-breeding area with severe phosphorus deficiency, high alkalinity and salinity levels, and periodic exposure to waterlogging [4]. There are few native legumes in these natural grasslands and none with significance as cattle fodder, so forage quality needs to be improved by introducing other legumes [5]. Under these constraining conditions, Lt is regarded as a “keystone species” for cattle nutrition [6] and could be used to improve the forage quality produced by natural or promoted grasslands [7]. However, even though Lc commercial varieties are less tolerant to these environments, they are planted over most of the sown area [1], probably because they contain foliar condensed tannins [6]. This is an important forage quality trait that contributes to ruminant diet, preventing cattle bloat, improving protein fraction assimilation, reducing intestinal parasites and reducing enteric fermentation, thereby decreasing greenhouse gas production [8]. Because Lt lacks condensed tannins, it has been crossed with a wild diploid Lc accession to produce fertile interspecific hybrids with adequate foliar condensed tannin levels inherited from Lt. To date, these interspecific hybrids have generally been found to have higher levels of tolerance to abiotic stresses than their parental accessions, particularly regarding long-term partial submergence stress [4] and saline conditions [9].

This region is also characterized by the presence of soil Na_2_CO_3_ and NaHCO_3_, the main sources of high soil alkalinity and therefore high pH. These factors can disrupt the balance of ions, cause micronutrient deficiency due to alteration of micronutrient availability in soil [10–12] and affect plant growth, development and survival [13]. These alterations are usually accompanied by symptoms of nutritional deficiency, particularly in young leaves, including interveinal chlorosis, which results from low concentrations of photosynthetic pigments, chlorophylls and carotenoids [14–13–15]. As a result, photosynthetic rate, light absorption and photosystem II (PSII) efficiencies are markedly reduced [16]. As an alternative to inorganic fertilization, plant growth-promoting bacteria (PGPB) can facilitate plant performance through a number of different bio-mechanisms such as adjustment in root size and morphology, modification of nitrogen accumulation and metabolism, and increased uptake of certain minerals [17,18]. In such regard, *Pantoea eucalypti* was isolated from alkaline soils from the Flooding Pampa and found to have the ability to solubilize phosphates and Fe, serving as a PGPB for Lt under both greenhouse [19] and outdoor conditions [20]. Furthermore, *P. eucalypti* proved to promote morphological, biochemical and molecular responses in *L. japonicus*, triggering the activation of “plant machinery” to optimize Fe metabolism and, as a result, improve the photosynthetic performance of Fe-inefficient plants under alkalinity [18].

The aim of the present work was to study Lt and Lc physiological responses to alkaline stress and to determine whether the interspecific hybridization of Lt × Lc, in addition to improving agronomic and qualitative traits of the *Lotus* genus, improves the ability to grow in marginal environments such as alkaline soils. The impact of inoculation with *P. eucalypti* was evaluated in all three accessions in the hope of developing strategies for improved legume nutrition under alkaline stress.

## MATERIALS AND METHODS

### Plant material and growth conditions

Seeds of a naturalized Argentine accession *Lotus tenuis* “Pampa INTA” (Lt), a diploid accession of *L. corniculatus* cv. “Charlii” (Lc), and an F1 population of *L. corniculatus* × *L. tenuis* (LtxLc) were used in this study*. L. corniculatus* seeds come from a wild population recovered from an area known as Devesa de El Saler in Spain, previously described by Escaray et al., 2014 [6]. Seeds were scarified with sulfuric acid (98%), washed in distilled water and sown in Petri plates containing water-agar (0.8%). Plates were incubated until the emergence of cotyledons in a growth chamber with a 16/8 h photoperiod (day/night) at 24/19°C, 60/80 ± 5% relative humidity under fluorescent light bulbs providing 200 μmol photons m^−2^ s^−1^. Seven-day-old seedlings were transferred from plates to plastic pots containing autoclaved sand and irrigated with half-strength Hoagland’s nutrient solution composed of 3 mM KNO_3_, 2 mM Ca(NO_3_)_2_·4H_2_O, 1 mM MgSO_4_·7H_2_O, 0.5 mM NH_4_H_2_PO_4_, 50 μM NaFeO·8EDTA·2H_2_O, 4.5 μM MnCl_2_·4H_2_O, 23 μM H_3_BO_3_, 0.16 μM CuSO_4_·5H_2_O, 17.5 μM ZnSO_4_·7H_2_O, and 12 μM Na_2_MoO_4_·2H_2_O [18]. After root development, each seedling of approximately one- month- old, was transferred to a beaker containing 150 mL of nutrient solution, where each treatment was assayed. The roots were protected from light by wrapping the beakers in black paper. The nutrient solution was replaced every 3 days.

### Bacterial strain and evaluation of endophyte colonization

A phosphate-solubilizing and siderophore-producing bacterium, *Pantoea eucalypti* M91, was used in this work [18,19]. The bacterial strain was isolated from the rhizosphere of *L. tenuis* growing in a typical lowland of the Salado River Basin [19]. The inoculant was prepared following Castagno et al., 2014 [20] and the inoculation treatment consisted of a bacterial suspension (1 × 10^8^ cells per mL) added to each beaker and refreshed whenever the nutrient solution was replaced.

In order to evaluate the endophytic bacterial colonization on the plants, *P. eucalypti* M91 was designed to express the *Red Fluorescent Protein* (RFP) as explained in Campestre et al., 2016 [18]. During harvest, plants were divided into roots, stems and leaves and analyzed by an epifluorescence microscope (Nikon Eclipse E600) equipped with a 450–490 nm excitation filter and a 520 nm long pass emission filter. Images were acquired with a Nikon DS-Qi1Mc video camera.

### Experimental design

The experiments followed a completely randomized design. All the accessions were evaluated under four conditions: i) Control, irrigated with nutrient solution for 3 weeks; ii) Alkalinity, irrigated with nutrient solution for 1 week, followed by the addition of NaHCO_3_ 10 mM to the nutrient solution for 2 weeks; iii) Control + Inoculation, control conditions plus the addition of the bacterial strain; iv) Alkalinity + Inoculation, alkalinity conditions plus the addition of the bacterial strain. The pH of irrigation solutions was monitored with a pH meter (HI 255, Hanna Instruments) and maintained at 6.2 and 8.2, for control and alkaline treatments, respectively. There were 8 beakers per treatment (each plant per beaker was considered as one biological repetition, n = 8).

### Determination of biomass

Tissues from harvested plants were divided into roots and shoots. Their dry matter (DW) was determined after drying at 70 °C until constant weight.

### Determination of H+ extrusion by roots

H^+^ extrusion was determined according to Campestre et al. (2016) [13] with some modifications. Root H^+^ extrusion was determined at 24 and 72 hours after the replacement of the hydroponic solution of plant cultures from all the assays during the last week of treatment, using a digital pHmeter. For each treatment, a control beaker without a plant was used. The pH of the hydroponic solution, pH 6.2 and 8.2 for control and alkaline plants, respectively, was only adjusted at the initial time without changing the volume of the solution throughout the assay (3 days). The pH gradient value (ΔpH) was calculated as:

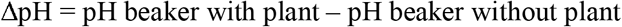

### Analytical determinations

Fe^2+^ was determined according to Campestre et al. (2016) [18]. Roots and shoots were harvested and carefully washed with deionized water. Plant organs were cut into small pieces with clean scissors. Fresh tissue samples (100 mg) of each organ were shaken in 1.2 mL of 80 mM 2.29-dipyridyl-HCl (pH 3.0) in 10% methanol in the dark (24 h). Extracts were passed through a 0.45 μm syringe filter and 1 mL of the filtrate volume was assayed at 522 nm using a spectrophotometer (Zel-tec ZL-5000 UV/VIS Spectrometer, Argentina). Fe^2+^ concentrations were calculated from a standard curve using a Fe atomic absorption standard solution.

Nine other elements (P, Mg, K, Ca, Fe, Zn, Cu, Mn, Mo) were determined as Dharmendra (2015) [21] with some modifications. 100 mg of dry material from roots and shoots were placed in glass vials and reduced to ashes at 550°C for 48 h. The ashes were digested with 3 mL of 65% HNO_3_ + 0.5 mL of H_2_O_2_. Extracts were filtered and diluted to a final volume of 10 mL with deionized water. Elemental analyses were performed using MP-AES (Agilent 4200, Agilent Technologies).

The analytical wavelength (nm) set for each analyte was: P (214.915); Mg (285.213); K (766.491); Ca (422.673); Fe (371.993); Zn (213.857); Cu (324.754); Mn (403.076); Mo (379.825).

### Determination of chlorophyll fluorescence fast-transient test

Non-invasive chlorophyll fluorescence fast-transient test (JIP test) was performed according to Gazquez et al. (2015) [22] on blades of the second expanded leaf with a portable chlorophyll fluorometer (Pocket PEA, Hansatech Instruments, UK). The parameters analyzed by JIP were maximum quantum yield of primary PSII photochemistry (F_v_/F_m_), specific energy flux of absorbed photon per PSII RC (ABS/RC), specific energy flux of dissipated excitation energy at time zero per PSII RC (DI_o_/RC) and performance index for energy conservation from photons absorbed by PSII antenna, to the reduction of Quinone b (PI_ABS_). For this purpose, leaves were covered with leaf clips to adapt them to darkness for 20 min. Then leaf clips were opened and samples were exposed for 3 s to 3500 μmol photons m^−2^ s^−1^ (637 nm peak wavelength). The pocket PEA software (PEA plus v1.1, Hansatech Instruments Ltd., UK) was used to analyze PSII properties according to Strasser et al. (2000) [23].

### NRAMP1 Gene expression

Total RNA was extracted from frozen roots using a Spectrum Plant Total RNA kit (Sigma) according to the manufacturer’s instructions. RNA quality was checked by agarose gel electrophoresis. After this, a DNAase treatment with an Ambion TURBO DNA-free kit was performed. Samples were quantified using a Synergy H1 Multi-Mode Reader, enabling the isolation of 2 μg of RNA for reverse transcription and cDNA synthesis. Quantitative RT-PCR of Strategy I NRAMP1 gene was performed. In order to conduct the analysis, 2.5 mL from a ten-fold dilution of the cDNA stock was further diluted to 15 mL with the primer mix (300 nM final concentration), 7.5 mL of Fast Start Universal SYBRH Green Master (Rox), and the required amount of double distilled water. Reactions were performed in an Mx3005P qPCR System with the help of the MxPro qPCR Software 4.0 (Stratagene, La Jolla, CA, USA). *EF* was used as the endogenous control (ID: AY633710). The following primers were used: NRAMP1 forward 5’-CAG AGA TTT GGG GTG AGG AA-3’, NRAMP1 reverse 5’-TGG TCT CGT CTC ACC TCC TT-3’. Results were expressed relative to the expression level of the control inoculated plants as relative units (RU), after being normalized to *EF* expression.

### Statistical analysis

Data were subjected to a two-way analysis of variance and Duncan’s test was used for multiple comparisons (*P* <0.05). The Shapiro-Wilk test to determine normality and Levene’s test to assess the equality of variances were used. All measurements were performed on 8 plants (= 8 biological replicates). All statistical procedures were carried out using the INFOSTAT statistical package (Version 2010, InfoStat group, FCA, Universidad Nacional de Córdoba, Argentina).

## RESULTS

### Plant growth and phenotype

The results revealed biomass variations in *Lotus* accessions regarding the response to alkaline stress and inoculation (Fig. 1). While alkaline treatment caused a considerable reduction in Lc total plant biomass of about 40%, Lt and LtxLc total biomass did not show any negative response to alkalinity. With respect to inoculation effect on total plant biomass, Lc increased its dry weight by about 50% under alkalinity whereas Lt had a positive response under control conditions of about 40%. In contrast, LtxLc did not show any differences between treatments. These responses occurred at shoot biomass level but not at root biomass level, with shoot biomass level displaying a pattern of response similar to that of total plant biomass. With regard to root:shoot biomass partitioning, only Lc showed differences in favor of shoot biomass under the control-inoculated treatment. In summary, the negative effect of alkalinity was particularly observed in the shoot biomass of Lc after 10 days of treatment (Fig. 2c) and in the development of interveinal chlorosis in the youngest leaves of Lc after 15 days of treatment (Fig. 2e). Moreover, favorable effects of Lc inoculation were observed as an increase in shoot biomass (Fig. 2d) and greening of young leaves (Fig. 2f). Additionally, all accessions were tested for endophytic colonization. The *P. eucalypti* strain harboring *Red Fluorescent Protein* (RFP) expression was found as an endophyte in the leaves, stems and roots of all three accessions under the four treatments assayed (Fig. S1).

**Figure 1.**
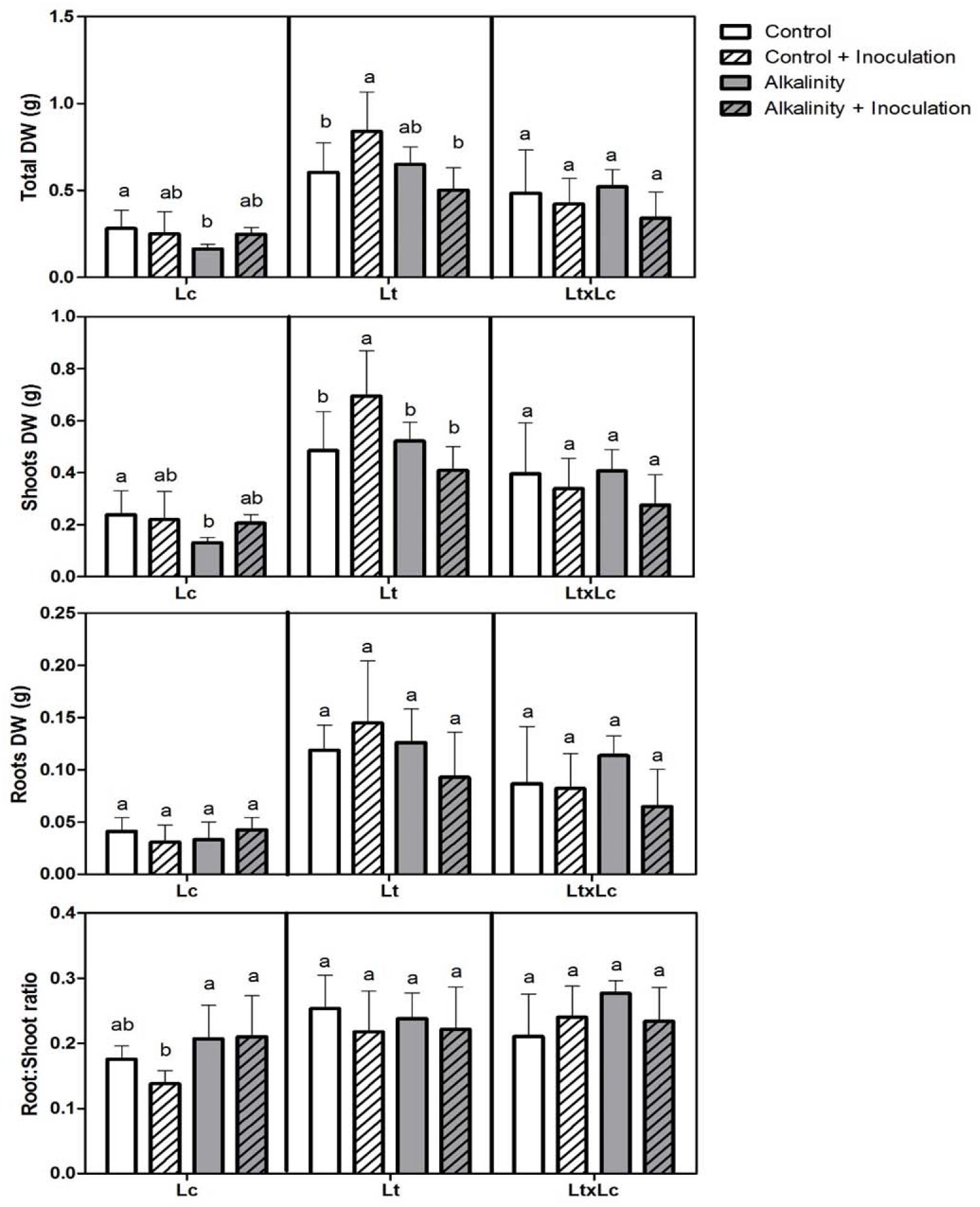
*Biomass production during control and alkaline treatments. Response to inoculation with a P. eucalypti strain.* Lc: *Lotus corniculatus*, Lt: *Lotus tenuis*, LtxLc: interspecific hybrid *Lotus tenuis* × *Lotus corniculatus.* Determination of total (a), shoot (b) and root (c) dry weight (DW) (g) and Root:shoot partitioning ratio. Bars (mean ± SD; n = 8) with different letters represent significant differences between treatments (Duncan’s test; P◻ 0.05).

**Figure 2.**
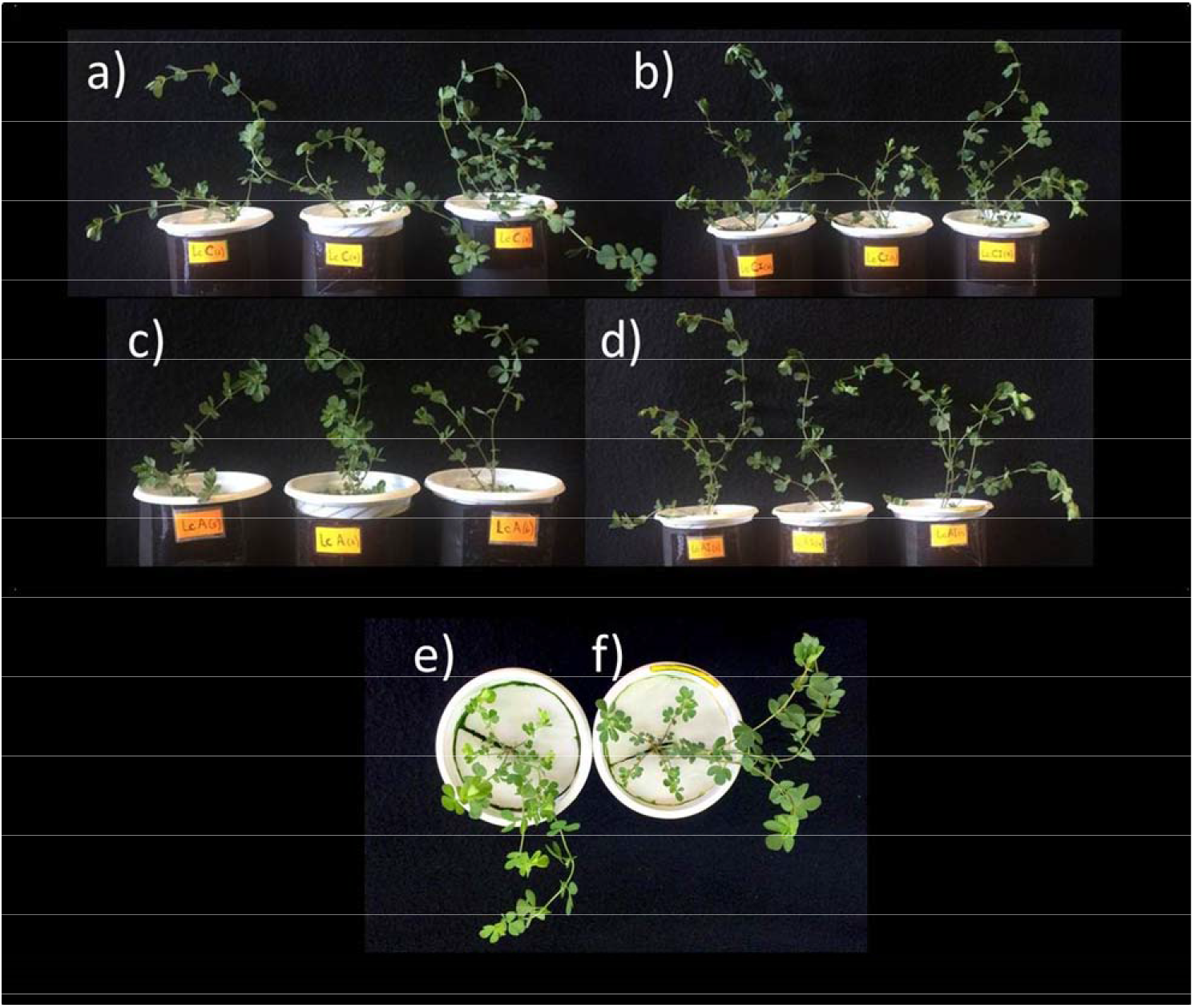
Phenotypic observation of Lc (*Lotus corniculatus*) plants under 10 days of exposure to control (a), control-inoculation (b), alkaline (c) and alkaline-inoculation (d) treatments. Interveinal chlorosis in young leaves of Lc plants (e) and greening of the leaves from inoculated Lc plants (f) after 15 days’ exposure to alkalinity.

### Evaluation of root H^+^ extrusion

Root H^+^ extrusion was determined at 24 and 72 h after the replacement of the hydroponic plant culture solution during the last week of treatment (Fig. 3), coinciding with the beginning of interveinal chlorosis in apical leaves of alkaline treated plants. At 24 h, all the evaluated accessions demonstrated a pronounced reduction in pH under control treatment and an increase in pH under alkalinity. Meanwhile, plant inoculation tended to bring the pH to its baseline value, increasing the pH under control treatment and decreasing it under alkalinity. On the other hand, at 72 h, Lc and LtxLc showed a similar pH response to that obtained at 24 h under control and alkaline treatment. In contrast, Lt showed no difference between treatments. Regarding the response to inoculation, only Lc showed differences in the control condition, while no difference was obtained with alkalinity for any of the three accessions. However, Lt and LtxLc pH under alkalinity seemed to be less alkaline at 72 h than at 24 h, while Lc seemed to maintain its gradient.

**Figure 3.**
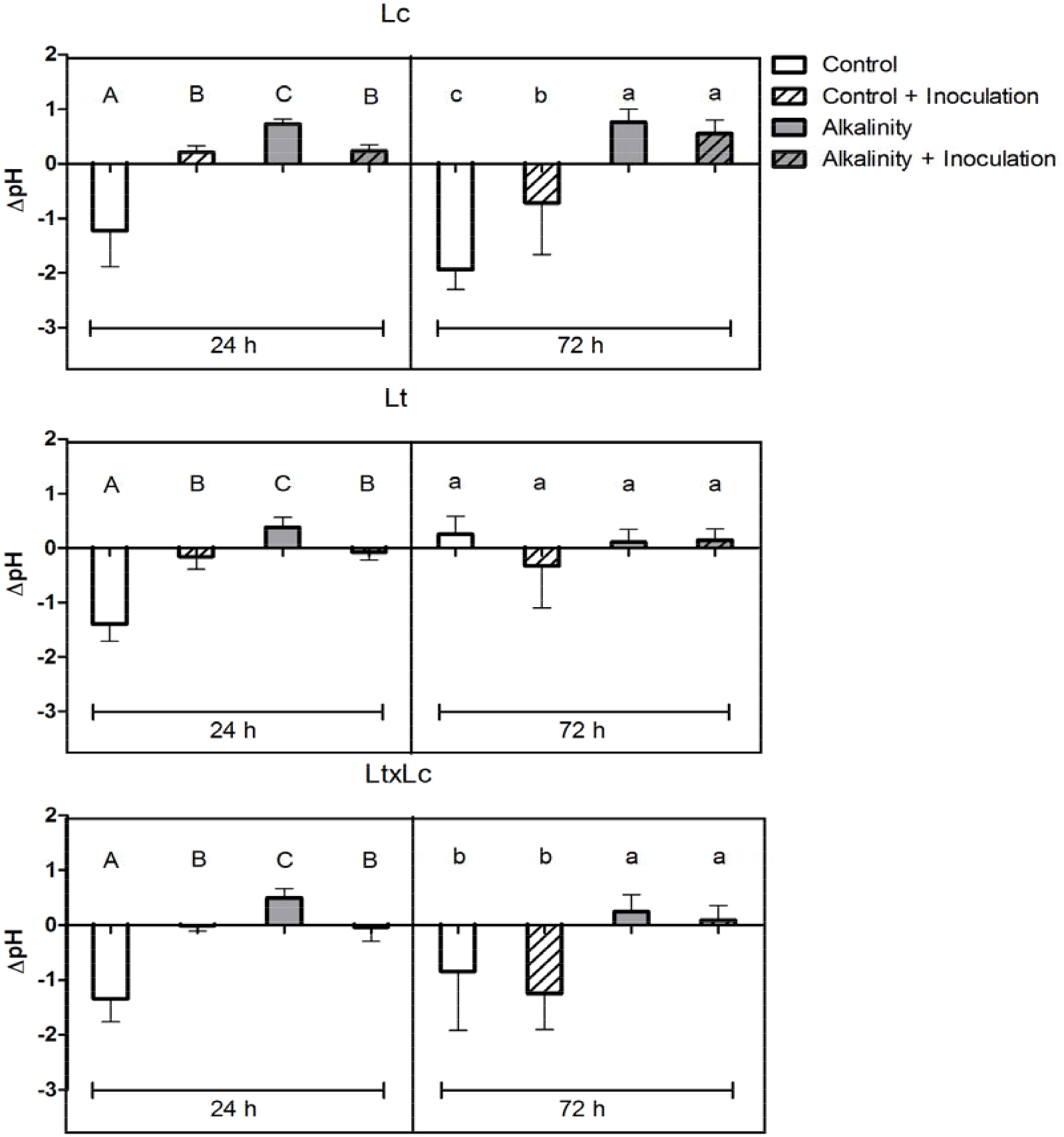
*H^+^ extrusion by roots of Lotus spp. plants grown under control, control-inoculation, alkalinity and alkalinity-inoculation treatment.* Lc: *Lotus corniculatus*, Lt: *Lotus tenuis*, LtxLc: interspecific hybrid *Lotus tenuis* × *Lotus corniculatus.* The pH of the nutrient solution was measured at 24 and 72 h and a control beaker without plant was performed for each treatment. ΔpH was calculated as: ΔpH = pH beaker with plant – pH beaker without plant. Points (mean ± SD; n = 8) with different letters (Capital for 24 h and lowercase for 72 h) represent significant differences between treatments (Duncan’s test; P◻ 0.05).

### Determination of selected micro and macronutrients in shoots and roots of *Lotus* spp. plants

As mentioned above, interveinal chlorosis was only observed in the youngest leaves of Lc during the last week of alkaline treatment (Fig. 2e). The Fe^2+^ determination indicated a drop in shoot Fe^2+^ content in Lc (around 70%) and LtxLc (about 37%) under alkalinity with respect to the control (Fig. 4). However, Lt increased its shoot Fe^2+^ content to about 90% under the same treatment. Fe^2+^ content was higher in the alkalinized roots of all three accessions than in the control treatment; however, Fe^2+^ accumulation was higher in Lt and Lc (4-fold and 3-fold, respectively) than in LtxLc (where it was 1.5-fold higher). In contrast, inoculation caused a considerable increase in shoot Fe^2+^ content in Lc, Lt and LtxLc under alkaline treatment, accompanied with a decline in root Fe^2+^ content in Lc under the same treatment.

**Figure 4.**
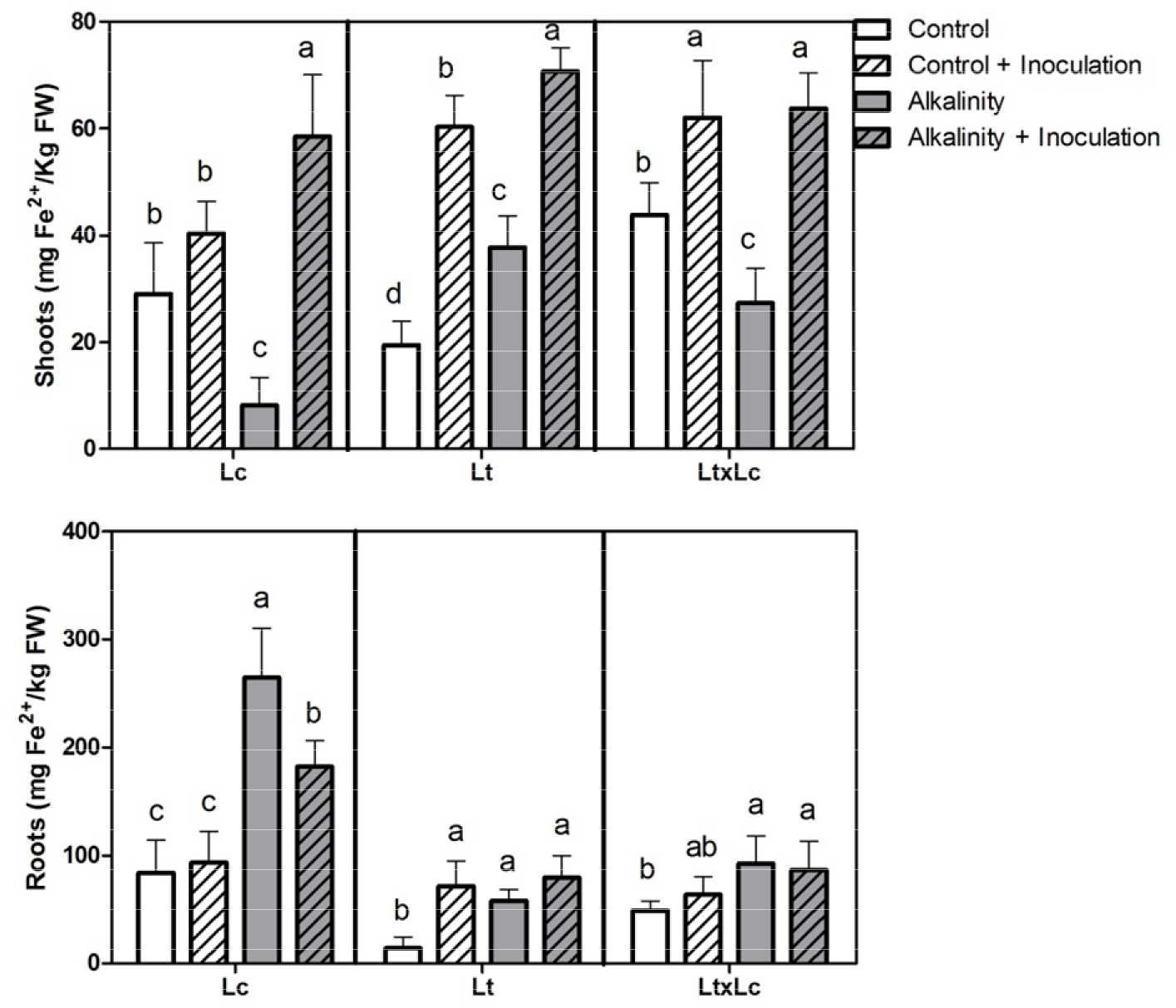
*Fe^+2^ content in Lotus spp. plants under control, control-inoculation, alkalinity and alkalinity-inoculation treatment*. Fe^+2^ content (mg/Kg FW) in shoots and roots were determined by spectrophotometry. Lc: *Lotus corniculatus*, Lt: *Lotus tenuis*, LtxLc: interespecific hybrid *Lotus tenuis* × *Lotus corniculatus.* Bars (mean ± SD; n = 8) with different letters represent significant differences between treatments (Duncan’s test; P◻ 0.05).

Despite these results, total shoot Fe content did not differ in Lt and LtxLc for any of the four treatments (Fig. S2). Differences were only observed in Lc, with a 2-fold decrease between control and alkaline treatment and a 2-fold increase due to inoculation under alkalinity. Other microelements, which are generally affected in plants exposed to alkaline stress, were also determined (Zn, Mn, Cu and Mo) (Fig. S2). Our results showed that the shoot content of each element varies among accessions, but in general, Zn was negatively affected in all three accessions, Cu in both parental accessions, and Mo in LtxLc when exposed to alkalinity. On the other hand, Mn content showed no difference in Lc, but an increase in Lt and LtxLc between control and alkaline treatment. As a consequence of inoculation, Zn content increased in Lt, Mn and Cu content were enhanced in Lc and no effect was detected in any of the accessions regarding Mo content under alkaline treatment. In addition, the macronutrients P, Ca and Mg were analyzed. P shoot content was notably affected in Lc and Lt under alkaline treatment. In contrast, Mg shoot content was not negatively affected in any of the accessions under alkalinity. Regarding Ca shoot content, only LtxLc increased its value under alkalinity. On the other hand, a similar decrease in P levels was observed in the roots in all three accessions (Fig. S3). Root concentrations of Fe only increased in Lc in plants that were inoculated under growth control conditions and subjected to alkalinity, in which inoculation caused a sharp decrease in total Fe levels, coinciding with the increase in Fe^2+^ in the aerial part (Fig. 3) and a reversal of the chlorotic effects caused by alkalinity (Fig. 2 e, f). Simultaneously, the levels of Mg^2+^ in the root increased under alkalinity and inoculation treatments in Lc and LtxLc, remaining unchanged in Lt. As was observed in shoots, the root levels of each element vary among accessions, but in general, Zn was diminished by inoculation in the hybrid and the opposite effect was observed in Lc subject to alkalinity. Cu levels remained unaltered in all three accessions, and Mo levels were reduced in Lc when exposed to alkalinity. Mn content increased by inoculated-alkaline treatment in all genotypes but only in Lt and LtxLc by alkaline stress.

### Influence of alkaline stress and inoculation on the PSII performance of *Lotus* spp. plants

PSII performance was estimated through the JIP parameters F_v_/F_m_, PI_ABS_, ABS/RC and DI_o_/RC (Fig. 5). The F_v_/F_m_ value was only reduced in Lc when it was cultivated under alkaline treatment (about 0.79) and the same effect was observed for PI_ABS_ (about 3.1). Interestingly, Lc plants cultivated under alkalinity and inoculated recovered F_v_/F_m_ and PI_ABS_ values equivalent to those obtained under control treatment, reaching about 0.82 and 4.8, respectively. In Lt, the ABS/RC ratio decreased under alkalinity compared to the control treatment and improved slightly with inoculation, while no difference was observed for the DI_o_/RC ratio between treatments. However, despite the fact that Lc did not show any differences in ABS/RC ratio, it dissipated more energy under alkalinity than the control plants. LtxLc showed no difference regarding energy absorbed but, in contrast to the other accessions, it dissipated less energy under alkalinity than the control plants.

**Figure 5.**
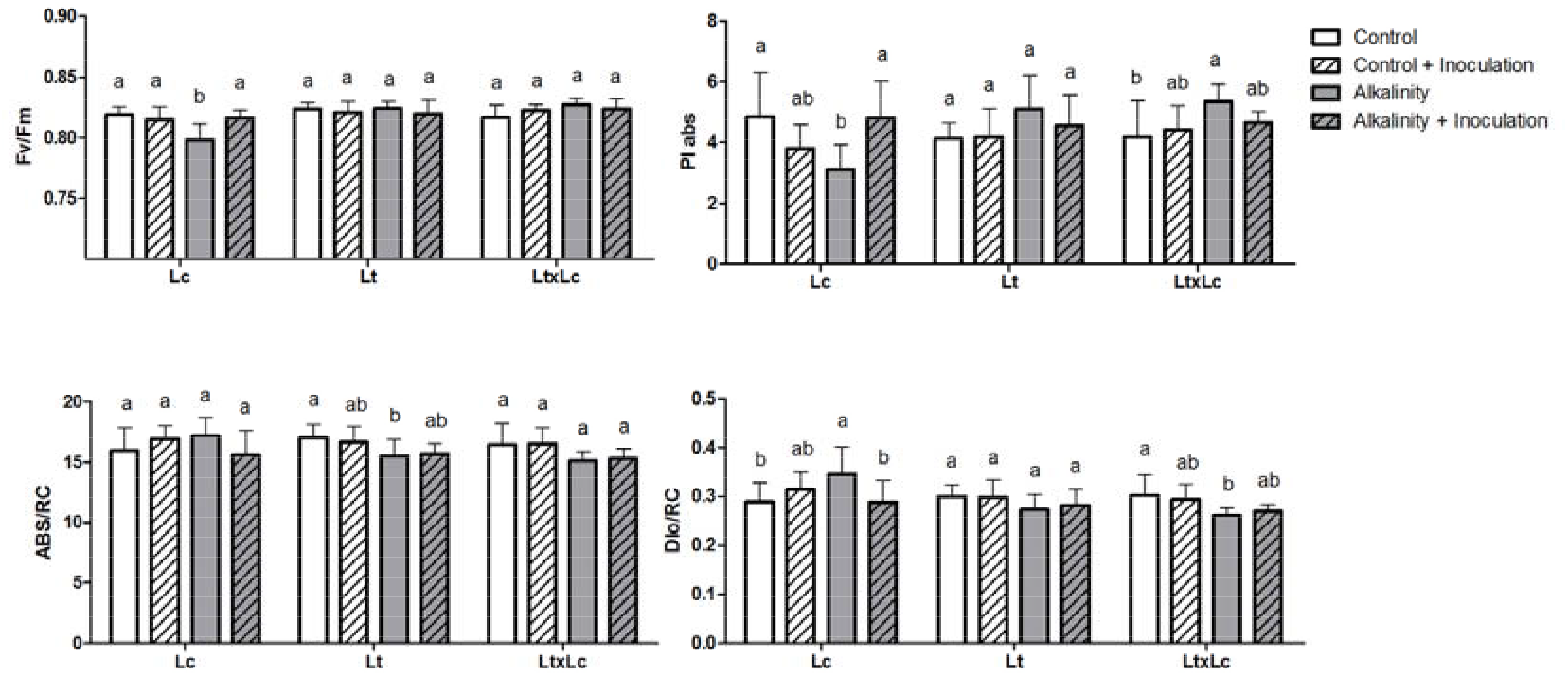
*JIP test in Lotus spp. plants exposed to control, control-inoculation, alkalinity and alkalinity-inoculation treatments. JIP* parameters tested were: Fv/Fm (a), PI abs (b), ABS/RC (c) and DIo/RC (d). Lc: *Lotus corniculatus*, Lt: *Lotus tenuis*, LtxLc: interspecific hybrid *Lotus tenuis* × *Lotus corniculatus.* Bars (mean ± SD; n = 8) with different letters represent significant differences between treatments (Duncan’s test; P◻ 0.05).

### NRAMP1 expression is induced in inoculated Lc roots under alkaline stress

The relative expression of the gene encoding the metal transporter NRAMP1 in roots of the three *Lotus* accessions was determined by RT-PCR (Fig. 6). Medium alkalization causes an increase in the relative expression of the NRAMP1 gene in Lc. The increase in its expression doubles in Lc with inoculation, a process that coincides with the reversal of chlorosis caused by alkaline stress (Figs.2e, f; 6).

**Figure 6.**
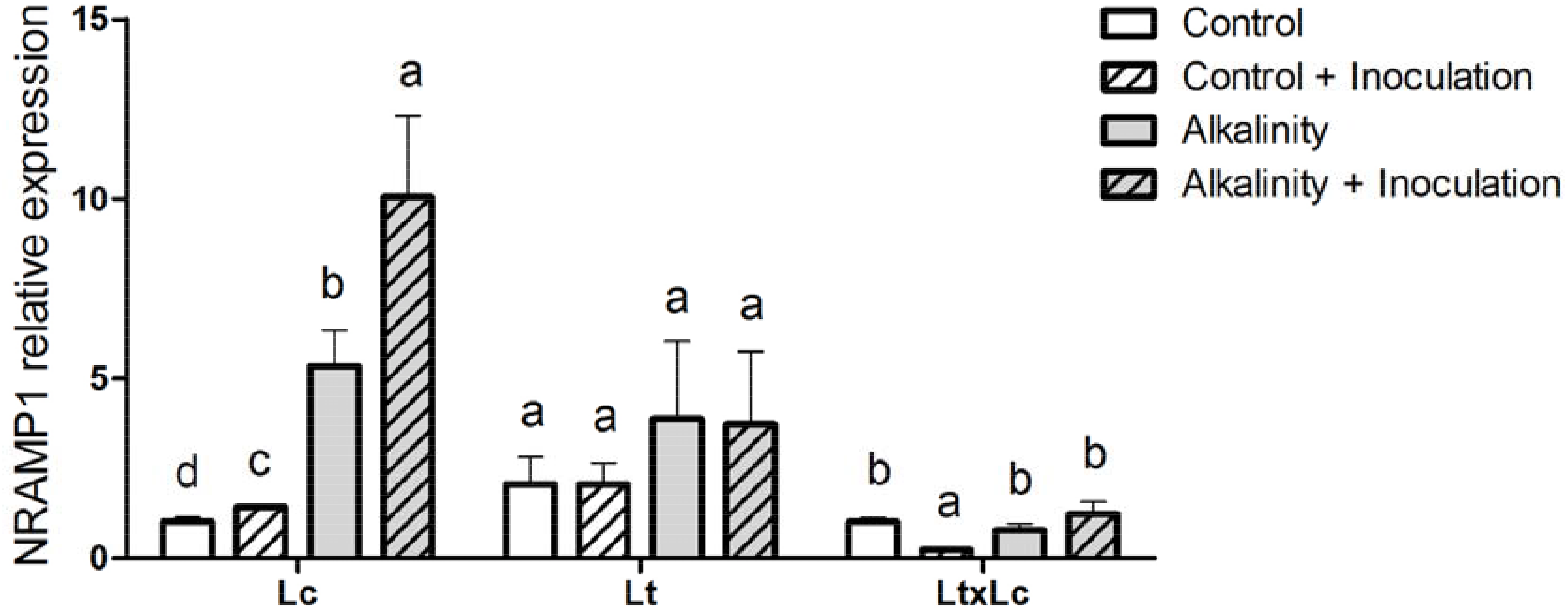
*Relative expression of* NRAMP1 *gene in parental and F1 hybrid roots exposed to control, control-inoculation, alkalinity and alkalinity-inoculation treatment.* Lc: *Lotus corniculatus*, Lt: *Lotus tenuis*, LtxLc: interspecific hybrid *Lotus tenuis* × *Lotus corniculatus.* Expression was determined by real-time RT-PCR analysis using EF-1α as the housekeeping gene. Bars (mean ± SD; n = 8) with different letters represent significant differences between treatments (Duncan’s test; P◻ 0.05).

## 4. DISCUSSION

Climate change is rapidly altering the stability of ecosystems and subjecting plants to complex eco-productive situations that include soil degradation and the coexistence of abiotic stresses [24]. In the Salado River Basin, Argentina, the proportion of cropland has increased in areas historically used for raising cattle [25]. The increasing use of land for agriculture has displaced cattle production to flooded lowland areas with saline-alkaline soils, which are restrictive for growing crops and forage species, including *Medicago sativa*, *Trifolium spp.* and other legumes. At the same time, soil degradation in many agricultural areas has led to a new concept involving livestock grazing as a way to make use of such areas in view of restoring them and improving farmers’ income. This technological proposal, however, has to take into account that plant growth, and therefore forage production, are affected by saline and alkaline stresses. It is thus essential to breed legumes with improved forage quality and performance under restrictive soil conditions in order to supply the growing demand for agri-food produced under environmentally-friendly conditions. Different strategies to improve legume forage quality are currently being developed in environments such as the Flooding Pampa [5, 26]. Some of these strategies involve identifying and selecting plant species that are adapted to highly constrained growing conditions. In this context, the biotechnological utilization of *Lotus* species is highlighted, based on their recognized tolerance performances against different abiotic stresses, high plasticity and nutritional value [5]. Currently, different *Lotus* species play important roles in the forage supply in South America, Australia and Europe. In particular, tetraploid Lc varieties have been extensively used due to their moderate level of proanthocyanidins (PA), which contributes positively to ruminant diet, increasing protein fraction assimilation, preventing cattle bloat and reducing intestinal parasites [6]. In contrast, Lt populations do not have significant levels of these secondary metabolites. Nevertheless, *Lotus* has become successfully naturalized in the Flooding Pampa, which is why some ecologists refer to it as a Keystone species. In addition, the tetraploid commercial cultivars of Lc have failed in the same ecosystem due to their susceptibility to the abiotic stress conditions of constrained soils where Lt occupies an environment characterized by the absence of native legumes with adequate forage productivity. Lt forage quality is comparable to that of other forage legumes such as *Medicago spp*. or *Trifolium spp*. However, its PA levels in leaves are low, decreasing its nutrient quality traits. In addition, Lt and Lc are considered salt-tolerant glycophytes, but the adaptability of tetraploid Lc (and some populations of Lt obtained through breeding) to constrained environments has been largely affected in commercial cultivars because breeding programs focus on dry matter production under nonrestrictive conditions. Based on this observation, a diploid Lc accession has been collected from the Valencia Albufera in Spain in an alkaline-salty area which is periodically flooded by the sea, with the intention of incorporating new traits of abiotic stress tolerance and increasing nutritional values of *Lotus* cultivars through inter-specific hybridization. We hypothesize that this accession would show better adaptation to combined stresses of lowlands in the Salado River basin than commercial tetraploid Lc. It is frequently recognized that high variability in the germplasm used in forage breeding programs could be an advantage in the search for tolerant traits and the acquisition of new nutritional attributes. Although this new germplasm was tested under waterlogging stress [4] and salinity [9], its stress response has not been evaluated previously under alkalinity. Moreover, the role of a plant growth-promoting bacterial endophyte (PGPBE) on alkaline stress mitigation has also been assayed.

Although shoot biomass production under alkalinity has been used in traditional breeding programs as a single trait to select tolerant genotypes, its success has been limited because other traits associated with alkaline tolerance should be considered. In legumes, the main mechanisms involved in long-term persistence in alkaline soils remain unknown. *Lotus* spp. materials evaluated showed differential shoot and root biomass accumulation under growth control conditions (Fig. 1). These results are consistent with those observed for the same materials in previous studies [4,6,9].

As the average pH at the seawater surface is about 8.2 and has remained steady for millions of years, taking into account the origin of the native population of the Albufera de Valencia, we expected it to have high tolerance to alkalinity. However, alkaline treatment caused a considerable reduction in Lc total plant biomass contrasting with that observed in Lt and the interspecific hybrid. It was also the genotype with the greatest development of senescence processes under alkaline stress (Fig. 2c). This phenotype was reverted by inoculation with *P. eucalypti* (Fig. 2d, e, f), suggesting that the endophyte activates response mechanisms that improve its performance and alkaline tolerance. Lt was the only accession in which aerial biomass increased by inoculation under control growth conditions. Moreover, regarding root:shoot biomass partitioning, only Lc showed differences in favor of root biomass under the alkaline and control-inoculated treatments. Further studies are required to clarify the reasons that account for these two features.

Root proton-secretion via the activity of plasma membrane H+-ATPase is considered to be an adaptation of plants to alkaline stress [18, 27]. The present study found that, under alkaline stress conditions, inoculation with the endophyte causes acidification of the medium. It is a known fact that alkaline and Fe deficiency stresses highly up-regulated H+-ATPase-encoding genes in the roots [27, 28, 29]. The effect observed with inoculation suggests that the root plants under stress might be able to release more H+ to acidify the rhizosphere and maintain root elongation in Lc. This observation would justify the root/shoot ratio observed in Lc plants inoculated under alkalinity (Fig 1). In addition, acidification of the rhizosphere may be beneficial for acquirement of Fe, thereby improving *Lotus* spp. growth under high pH conditions. As has often been reported, under high pH conditions, the availability of Fe is quite limited for plant use due to precipitation of Fe [10, 13, 15, 18]. It has been well documented that higher plants use two major Fe uptake strategies (Strategies I; Fe reduction and Strategies II; Fe chelation) to acquire more Fe under this stress condition [13, 18]. In gramineous plants, Fe acquisition has been operated by only Strategy II, which can release mugineic acid family phytosiderophores to uptake Fe^3+^ from alkaline soils.

Previous reports [18] demonstrated that inoculation of *L. japonicus* ecotype Gifu with *P. eucalypt*i M91 under alkaline conditions reversed the development of interveinal chlorosis of the apical leaves due to low mobilization of Fe^2+^. In addition, it was observed that *P. eucalypti* M91 caused plant growth-promoting effect on the model *Lotus* species, producing pyoverdine-like and pyochelin-like siderophores under alkaline stress. Also, the PGPB strain alters root morphology, resulting in a herringbone pattern of root branching [10, 12, 18]. Additional features include improvement in Fe^2+^ transport to the shoots, acidification of the hydroponic solution of the plant cultures, and an accompanying increase in the efficiency of the PSII parameters. The plant alkaline stress mitigation was accompanied by an increase in the expression of the FRO1 and IRT1 genes and a significant increase in FRO activity. All these results are in line with the results obtained with the Lc accession, showing that *P. eucalypti* M91 has a beneficial effect on the Fe acquisition machinery of Strategy I, suggesting its potential as an inoculant for legume crops cultivated in alkaline soils.

In addition, alkalinity may cause nutrient deficiencies or imbalances, and plants present different strategies to respond [10, 12, 15, 18, 30]. The analyses of macro and micronutrients in roots and aerial parts revealed variations that depended on the accession of *Lotus* spp. The explanation for each of such differences requires a particular study for each element. However, they enabled the establishment of a good correlation between total Fe levels in the plant, and the strategy of mitigating alkaline stress mediated by the endophytic bacteria. Likewise, a decrease in P levels was determined, which would be explained by the hydroponic study carried out. However, the ability to solubilize and improve P nutrition in crop and model *Lotus* plants has been reported previously [19, 20], supporting potential use in bioinoculant formulations.

PSII activity, estimated by chlorophyll fluorescence measurements, was altered in the Lc plants under alkaline stress, but not in the Lt and Lt × Lc accessions and the Fv/Fm was recovered by inoculation (Fig. 5). It has been reported previously that the ◻_PSII_ parameter decreased in Lt plants under salt stress, suggesting that this species could lose carbon fixation efficiency under this stressed condition [6]. At the same time, higher NPQ values were observed in the salt-exposed Lt plants. Interestingly, similar results were observed under alkalinity but for Lc. An anticipatory induction of NPQ has been described as a plant adaptive response to dissipate excess excitation energy as heat. This photoprotection mechanism helps to minimize ROS production, and consequently protects from oxidative stress. Since the Lc genotype used herein came from an extremely saline-alkaline and frequently flooded environment, and it is possible that its high tolerance to salt stress was due to anatomical adaptations developed to grow under these constrained conditions. Because Lt has also been considered a tolerant glycophyte, the genetic pools of both *Lotus* spp. are invaluable for improving abiotic stress tolerance in the genus. It has been demonstrated previously that the LtxLc hybrid plants outperform the traits of their parental accessions and present improved nutritional characteristics for their use as forage [4,6,9]. Interestingly, under alkaline stress, the hybrid plants also seemed to outperform their parental accessions because no senescence symptom was observed. Moreover, in previous studies it was observed that net photosynthetic rate under saturating irradiance (Asat) did not change for Lt and LtxLc at the end of flooding stress treatment (55 days), while it was considerably lower for tetraploid accession of Lc under the same condition compared to controls [4]. These data are consistent with the differences observed in biomass accumulation. Lower Asat values could be partially explained through the impairment of PSII. Alterations in the PIabs parameter were consistent with the alterations observed for Asat values in the different accessions [4]. These results, added to the observations of the current study, show that interspecific hybridization generates plant material that better copes with abiotic stress situations, reducing its photosynthetic performance to a lesser degree.

Finally, as reported previously, AtNRAMP1 regulates Fe homoeostasis in *Arabidopsis*. Moreover, it is known that when this gene is incorporated into yeasts, it enables Fe uptake. The gene appears to be expressed preferentially in Fe-deficient roots, indicating its role in Fe uptake and transport. It has been shown that overexpression of AtNRAMP1 leads to an increased resistance to toxic Fe level [31], indicating it may participate in Fe remobilization under Fe deficiency. Beyond functioning in Fe transport, NRAMP genes have been also been demonstrated to perform wide-ranging transport activities for divalent transition metals in plants, including Mn^2+^, Fe^2+^, Co^2+^, Ni^2+^, Cu^2+^ and Zn^2+^ [32].

In recent years, several NRAMP genes have been identified in legumes. For example, a peanut NRAMP gene, AhNRAMP1, is significantly induced by Fe deficiency in roots and leaves, and heterologous expression of AhNRAMP1 in tobacco leads to Fe accumulation in young leaves and tolerance to Fe deprivation [33]. Moreover, in the model legume *Medicago truncatula*, MtNRAMP1, it is mainly located in the plasma membrane, with expression levels highest in roots and nodules, suggesting it was the major transporter responsible for apoplastic Fe uptake in rhizobia-infected cells [34]. More recently, it was demonstrated that NRAMP1 plays a pivotal role in Fe transport by cooperating with IRT1 to take up Fe in roots under stress condition [35]. Studies of the expression of this gene in *Lotus* spp. has not been reported previously. The induction observed in the Lc collected from the Albufera de Valencia suggests its contribution to the response of stressed plants with low levels of available Fe and would account for the better homeostasis of Fe in the hybrid material.

## 5. CONCLUSION

Taking into account all the antecedents evaluating *Lotus* spp. (model and forage), which included the parents of the interspecific hybrid LtxLc subjected to the stresses that usually coexist in soils affected by alkalinity, salinity and flood, we can conclude that the hybrid plants performed better than their parental accessions. This could be explained, at least in part, by the combination of responses from both parental accessions displayed by the hybrid under abiotic stresses. The new gene assortments obtained by the interspecific cross between Lt and diploid Lc accession has led to the generation of novel genotypes with a high potential to be used as forage in the lowlands of Salado River Basin. However, further studies are needed to understand photosynthetic acclimation to the combined stress condition.

## ACKNOWLEDGEMENTS

This work was supported by grants from the *Agencia Nacional de Promoción Científica y Tecnológica* (ANPCYT-Argentina) /FONCyT-PICTs 1560, 1612, 3648 and 3718 and Consejo Nacional de Investigaciones Científicas y Técnicas (CONICET-Argentina)/PIP 0980. M.P.C., C.J.A were CONICET post-doctoral fellows. VGM is doctoral fellow of CONICET, whereas N.L.C and O.A.R. are members of the CONICET Researchers Career.

## AUTHORS’ CONTRIBUTIONS

M.P.C. designed and performed all the experiments, and analyzed data; N.L.C. performed some experiments and analyzed some data; C.J.A. and V.G.M performed some experiments and analyzed data; O.A.R. conceived the project and designed and supervised all the experiments. The article was written by M.P.C., C.J.A. and O.A.R.

## Supporting material

**Supplementary Figure 1.**
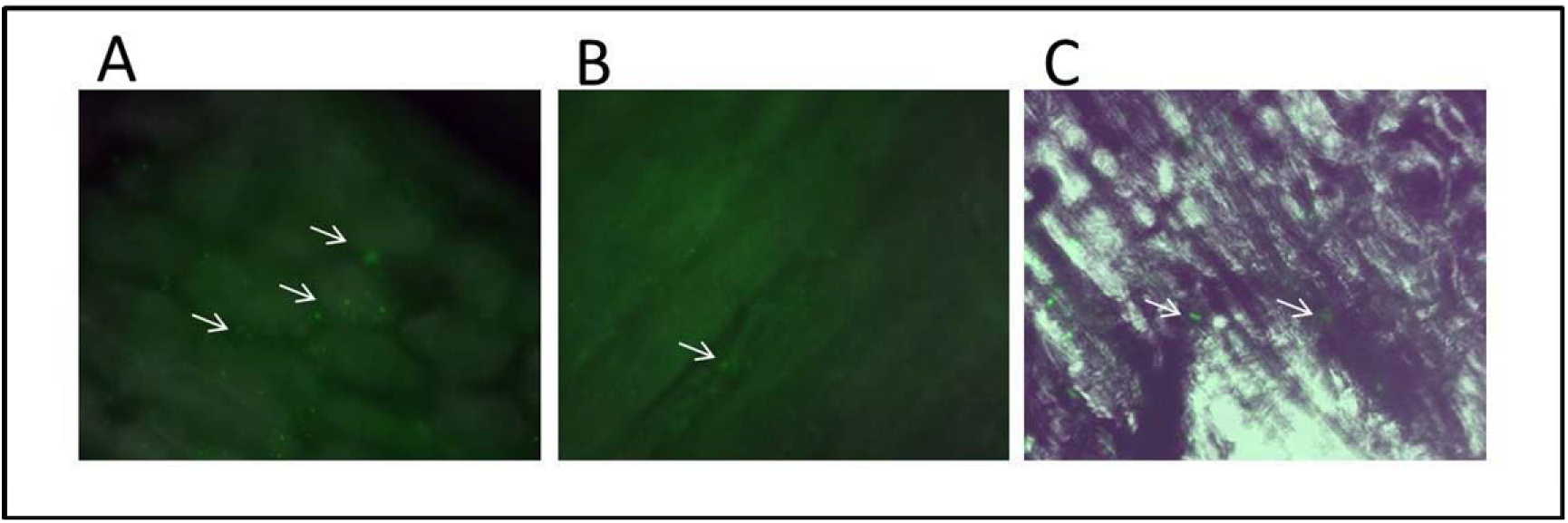
*Representative images of the endophytic colonization in the Lotus spp. plants visualized by epifluorescence. P. eucalypti* fluorescent cells (white arrows) are found colonizing intercellular spaces of (A) leaves, (B) stems and (C) roots of the inoculated plants (100X).

**Supplementary Figure 2.**
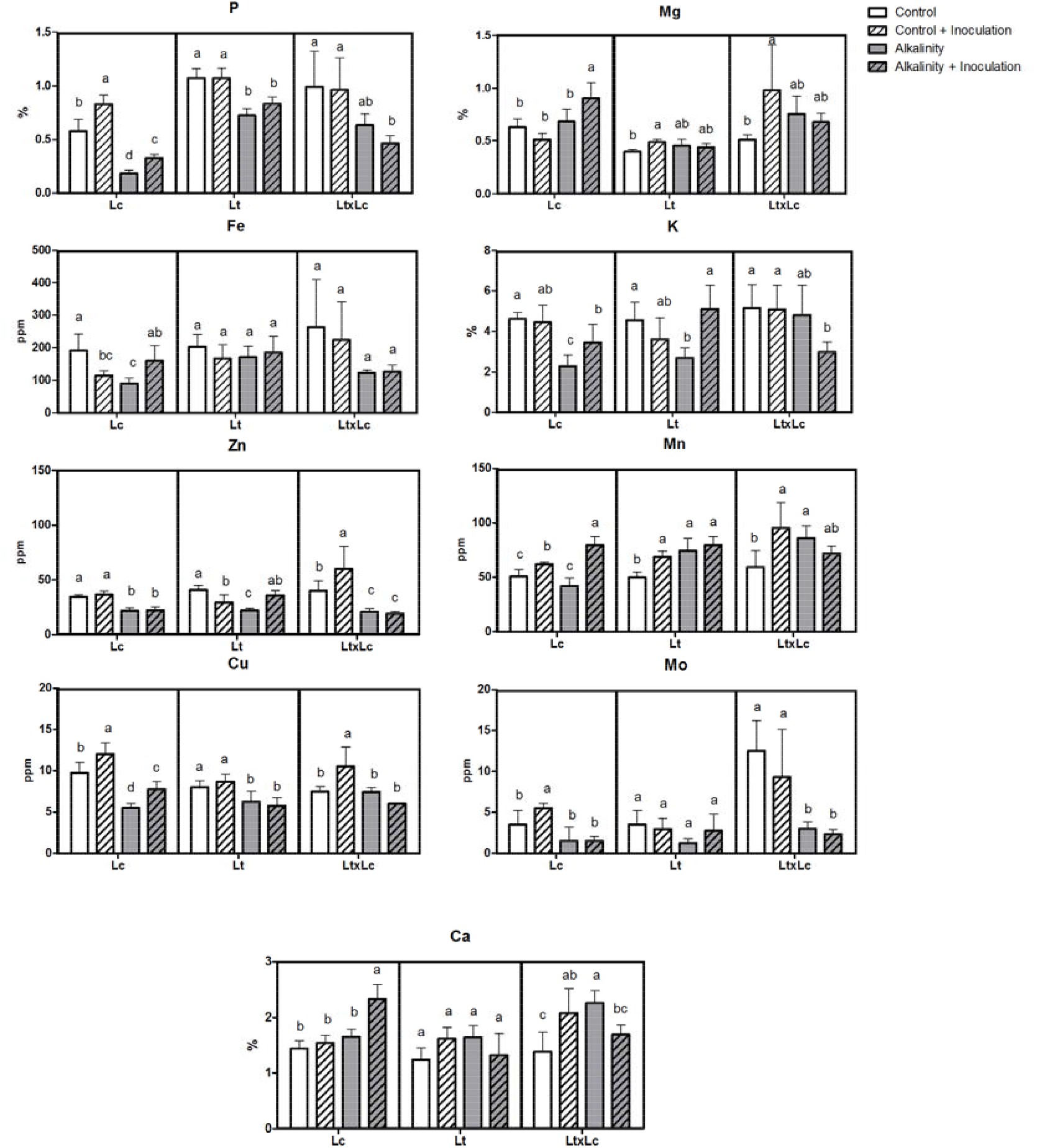
*Micro (Fe, Zn, Mn, Cu, Mo) and macronutrient (P, Ca, K and Mg) content changes in* shoots of *Lotus spp. plants under control, control-inoculation, alkalinity and alkalinity-inoculation treatment*. Micronutrients (ppm) and macronutrients (%) were measured by MP-AES. Lc: *Lotus corniculatus*, Lt: *Lotus tenuis*, LtxLc: interspecific hybrid *Lotus tenuis* × *Lotus corniculatus.* Bars (mean ± SD; n = 8) with different letters represent significant differences between treatments (Duncan’s test; P◻ 0.05).

**Supplementary Figure 3.**
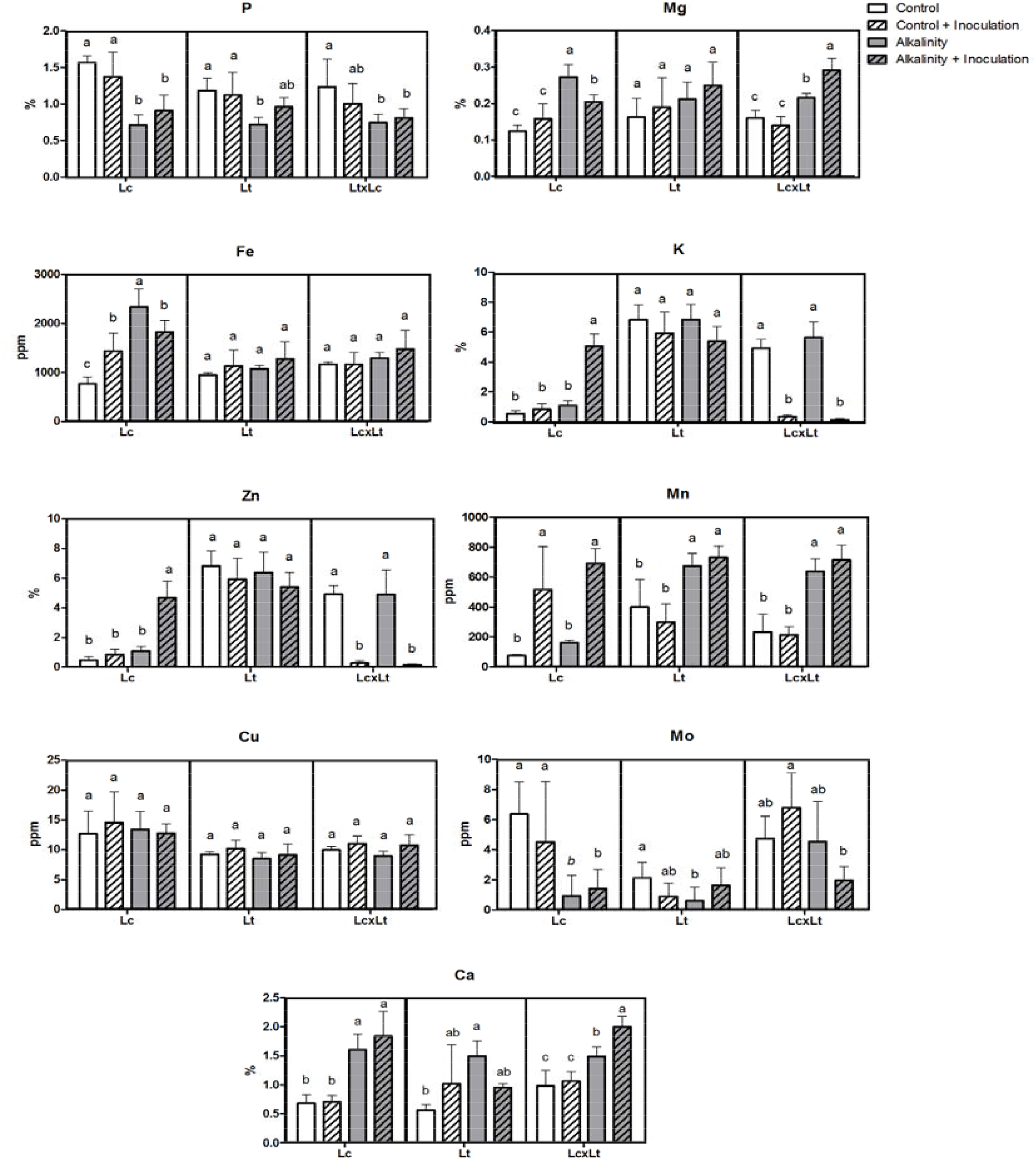
*Micro (Fe, Zn, Mn, Cu, Mo) and macronutrient (P, Ca, K and Mg) content changes in roots of Lotus spp. plants under control, control-inoculation, alkalinity and alkalinity-inoculation treatment*. Micronutrients (ppm) and macronutrients (%) were measured by MP-AES. Lc: *Lotus corniculatus*, Lt: *Lotus tenuis*, LtxLc: interspecific hybrid *Lotus tenuis* × *Lotus corniculatus.* Bars (mean ± SD; n = 8) with different letters represent significant differences between treatments (Duncan’s test; P◻ 0.05).

